# Functional consequences of fast-spiking interneurons in striatum

**DOI:** 10.1101/2024.09.17.613386

**Authors:** Arvind Kumar, Lihao Guo

## Abstract

The striatum features a distinct network characterized by a high degree of shared feedforward inhibition (FFI) from a mere 1% of fast-spiking interneurons (FSI). We investigate the potential roles of this extensively shared FFI in striatal function beyond inducing synchrony. Our findings reveal that FSIs increase the across-trial variability of striatal responses to cortical stimuli and, combined with recurrent inhibition, lead to a ‘correlation attractor’ of striatal activities, i.e., weakly correlated inputs result in more correlated responses and vice versa. Thus, we uncover a mechanism by which input correlation can be bidirectionally modulated, which is possible only because of high sharing of FSI inputs. We posit that the emergence of a correlation attractor leads to non-zero correlation level and variable rate trajectories of striatal responses across trials, hence beneficial for exploration in learning. However, given their role in across-trial variability, we argue that FSIs should be ‘disengaged’ from the MSNs during performance where stability across trials is required.

**Significance Statement:** Striatum is a network of inhibitory neurons. Fast spiking interneurons constitute about 1% of the striatal neural population and provide feedforward inhibition (FSI). Here, we unravel two novel ways in which FSIs may shape striatal function. Given the recurrent inhibition, it is assumed that striatum can only de-correlate inputs. We show that high sharing of FSI also renders the striatum an ability to correlate inputs. Thus, recurrent and shared FSI create a ‘correlation attractor’. Besides, we show that shared FSIs give rise to high across-trial variability. Therefore, we argue that FSIs are more crucial in learning as they provide the neural basis of exploration, but they may impair learned behavior due to high across-trial variability.

## Introduction

The striatum, the main input station of the basal ganglia, is a purely inhibitory network of medium spiny neurons (MSN) and several interneuron types. The MSNs which express either D1- or D2-type dopamine receptors form the bulk of the striatal network (≈ 95%) and interneurons such as cholinergic interneurons (ChIN), fast-spiking interneurons (FSI) and low-threshold spiking (LTS) neurons make the remaining 5% [Burke et al., 2017].

The FSIs form the classical feedforward inhibitory (FFI) motif in the striatum. That is, cortico-striatal inputs project onto both the MSNs and FSIs, with FSIs subsequently projecting onto the MSNs. Thus, MSNs receive cortical excitation (E) which is quickly followed by inhibition (I) from FSIs. FFI is one of the most ubiquitous motifs in the brain [see review by Isaacson and Scanziani, 2011], playing critical roles in defining the temporal window for spike discharge, modulating circuit gain [Isaacson and Scanziani, 2011], and selectively gating neural activity patterns [Kremkow et al., 2010]. Moreover, when synapses exhibit short-term facilitation/depression, FFI can modulate the dynamic range of neural output [Grangeray-Vilmint et al., 2018] and enhance the detection of signals with low signal-to-noise ratio [Tauffer and Kumar, 2021].

What sets the striatal FFI apart from other brain networks is the fact that only 1% of the striatal neurons are FSIs, yet the projection covers all MSNs. In fact, FSIs project onto MSNs with a connection probability over 60% within their axonal reach [Koos and Tepper, 1999; Planert et al., 2010; Cinotti and Humphries, 2022]. This is starkly different from the neocortex, cerebellum and hippocampus, where the FSI→pyramidal neuron (PYD) connection probability is between 10-20%. Such a high connection probability in a small FSI population implies that MSNs receive highly shared FFI, potentially introducing synchrony among MSNs [Yim et al., 2011; Gittis and Kreitzer, 2012].

Beyond synchronization, it remains unclear how FSIs may affect the striatal function, particularly the processing of cortical inputs which presumably carry multisensory information about the animal state and action-values [Wall et al., 2013]. Given the multisensory information, the basal ganglia is believed to do action-selection in a state-dependent and context-dependent manner. However, the classical go/no-go winner-take-all dynamics is not biologically plausible given the relatively weak recurrent inhibition between MSNs [Cui et al., 2013; Barbera et al., 2016; Klaus et al., 2017]. It is possible that heterogeneous responses of FSIs at millisecond-scale combined with the highly-shared FFI can regulate the signal transfer from the cortex to MSNs [Berke, 2011].

To understand how sharing of FFIs may affect the response of MSNs to cortical inputs, we used numerical simulations of a reduced striatal model comprising MSNs and FSIs (Figure 1 A). We systematically varied the number of FSIs in the model while keeping total FSIs input to MSNs constant. We found that besides introducing synchrony, FSIs affect the MSNs responses in two different ways: (1) Across trials, the FFI increases variability of striatal activities; (2) Within a trial, shared FFI and recurrent inhibition created a correlation transfer attractor such that when the striatum is driven by highly correlated (uncorrelated) inputs, ensembles of MSNs are de-correlated (correlated). These results suggest that FSIs may play a bigger role in the early phase of learning, when correlation structure from cortical inputs is less stable and high across-trial variability could be used for exploration.

**Figure 1:**
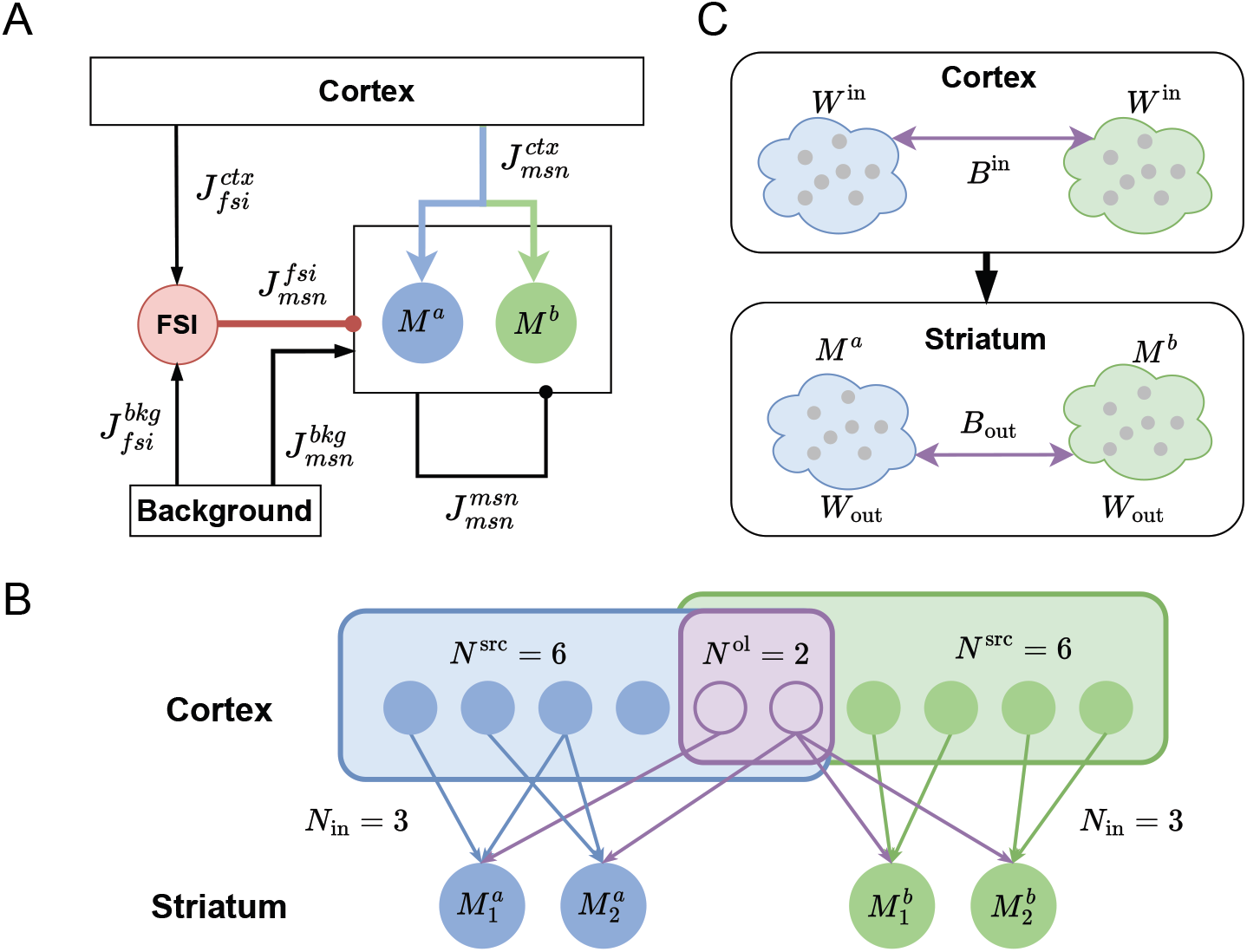
Simplified cortico-striatal circuit (**A**) The diagram of a simplified cortico-striatal circuit. Each arrow denotes connection from one population to another, where arrow-head is excitatory and circle-head is inhibitory. In the circuit, MSNs receive direct excitatory inputs from the cortex and relayed inhibitory inputs via FSIs. The spontaneous activities of the striatum are maintained by background inputs. The inhibitory projections from FSIs to MSNs 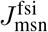 are much stronger than the recurrent connections between MSNs 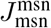 denoted by the arrow widths. (**B**) The feedforward connections from cortex to MSNs. Each node denote a neuron, either a pyramidal neuron in the cortex or an MSN in the striatum. The arrows denote the excitatory projections from cortex to MSNs. There are two MSN populations differentiated by the input source. The blue-colored MSNs receive input from the blue and purple cortical neurons, while the green-colored MSNs receive input from the green and purple cortical neurons. Each MSN neuron has a fixed input degree *N*_in_. (**C**) Correlation transfer from cortical input correlations to striatal output correlations. *W* denotes correlation within a group and *B* denotes correlation between groups.

Across-trial variability is a hallmark of neural activity and plays an important role in learning and behavior [Dinstein et al., 2015; Waschke et al., 2021]. Computationally, across-trial variability could be modulated by input structure [Litwin-Kumar and Doiron, 2012; Deco and Hugues, 2012], network connectivity [Bujan et al., 2015], or both [Guo and Kumar, 2023]. Our results here have uncovered that high sharing of FSIs could also be an equally important determinant of across-trial variability.

As our findings solely depend on the high degree of shared connectivity, implications of our results go beyond the striatum and apply to other rare cell types (i.e., those constituting less than 5% of the total population) found in neuronal networks in the brain. Thus, our findings also underscore the potential influence of rare cell types on behavior.

## Materials and Methods

### Experimental Design

For all experiments, the network structure is given as Figure 1 A with 2,500 MSNs. Because here we want to investigate the role of FSI (and its FFI), for a fair comparison we need to compare cortico-striatal network with and without FSIs as shown in Figure 1. To study the effect of the sharing of FSIs inputs by MSNs, we systematically varied the number of FSIs in our model (from 25 to 250) while keeping the number of FSI input per MSN constant as 15. Thus, we varied the sharing of FSIs input to MSNs from 60% to 6%. For each setting of the FSI population size and input parameters (see Input model) including the control case, we simulated 10 independent trials of the network activity for 2.5 s. The first 500 ms for each trial was for stabilization, and the remaining 2 s were used for analysis. Across the 10 independent trials, the cortical input was identical while the background noise was distinct with new realization for each trial.

### Neuron model

The neurons were realized as conductance-based leaky integrate-and-firing neuron [Meffin et al., 2004]. The subthreshold membrane potential dynamics of an MSN is given by the following equation:

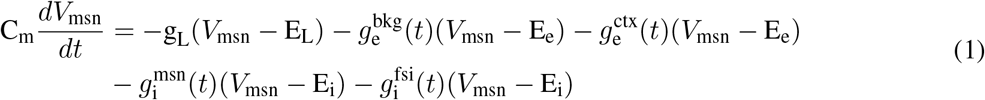

where *V*_msn_ (mV) is the subthreshold membrane potential, *C*_m_ is the membrane capacitance, *g*_L_ is the leak conductance and *E*_L_ is the resting membrane potential. When the *V*_msn_ reaches the threshold V_th_, a spike is elicited and the membrane potential is reset as *V*_msn_ = E_L_(mV) with a following refractory period of *t*_r_. The neuron is driven by conductance-based excitatory synaptic inputs (see Synapse model): 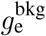 is the total background input, 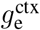 is the total excitatory input from the cortex, 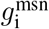 is the total inhibitory input from other MSNs and 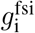 is the total inhibitory input from FSIs. The subthreshold membrane potential dynamics for FSI is similar, except without inhibitory input from FSIs. Each presynaptic spike was transferred as an alpha-function shaped conductance transient to the postsynaptic neuron. *E*_e_ and *E*_i_ are reversal potentials of the excitatory and inhibitory synapses, respectively. In the network, all striatal neurons were identical with parameters provided in the Table 1.

**Table 1:** Neuron parameters.

| Parameter<br>Symbol<br>Value | Membrane capacitance<br>$C_m$<br>250 pF | Leak conductance<br>$g_L$<br>16.67 nS | Resting potential<br>$E_L$<br>-70 mV | Spike threshold<br>$V_{\text{th}}$<br>-55 mV |
| --- | --- | --- | --- | --- |
| Refractory period<br>$t_r$<br>2 ms | Exc. reversal potential<br>$E_e$<br>0 mV | Inh. reversal potential<br>$E_i$<br>-85 mV | Rise time Exc. syn<br>$\tau_e$<br>0.2 ms | Rise time Inh. syn<br>$\tau_i$<br>15 ms |

### Synapse model

Neurons were connected using conductance-based synapses. Each incoming spike resulted in a conductance transient *g*_syn_(*t*) that decayed exponentially with a time constant *τ*_syn_:

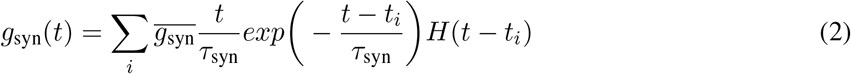

where *t*_*i*_ is the arrival time of *i*^*th*^ spike and *H* is the Heaviside step function. The maximal conductance for each synapse was given by 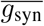. Time constants for excitatory and inhibitory synapses, i.e. *τ*_e_ and *τ*_i_, are given in the Table 1. The feedforward inhibition from FSI is dominated by GABA_A_ receptors [Tepper et al., 2008] with time constant in the range of 10ms, therefore, here we used a constant 15ms for *τ*_i_.

### Connectivity model

The MSNs were connected to each other independently with a connection probability of 10% (250 in-degree) and each MSN received 15 connections from FSIs. All connection strengths 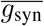 and synaptic delays are given in Table 2. The synaptic delay denotes the transmission time of a spike from a presynaptic neuron to a postsynaptic neuron. Consistent with experimental observations [Tepper et al., 2008], we assumed that the synaptic weight of feedforward inhibition from FSI is much stronger than the recurrent inhibition between MSN. We assumed that FSIs do not have any mutual connectivity via chemical or electrical synapses. Moreover, MSNs also did not project to FSIs.

**Table 2:**
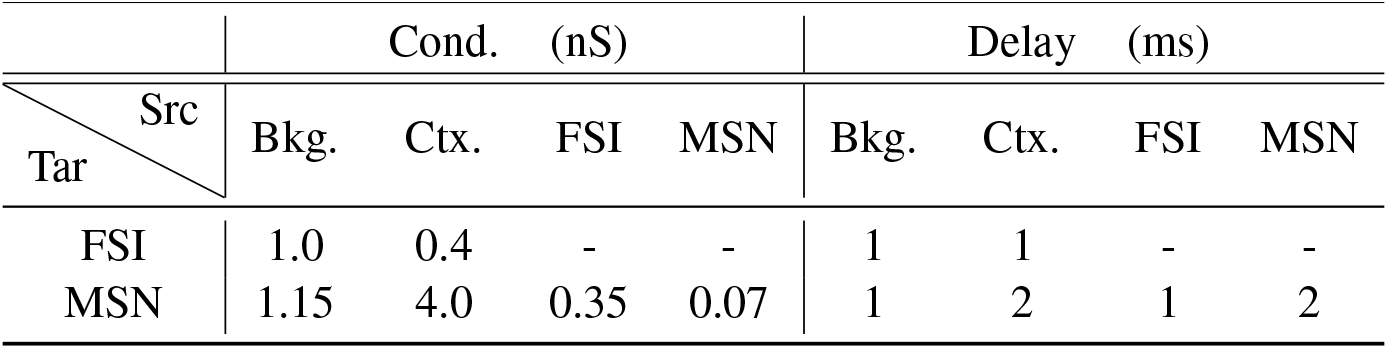
Network connectivity parameters.

### Background input

To mimic the ongoing activity, each neuron received excitatory inputs as uncorrelated homogeneous Poisson spike trains from the background input source with rates of 7.3kHz (4.8kHz for no FSI case to maintain a similar evoked firing rates). The background input sustains the spontaneous activities of MSN (around 1.5Hz) and FSI (around 5Hz). The synaptic weights of background inputs are given in Table 2.

### Input model

The two prominent features of *in vivo* spiking activities, weak pair-wise correlation and Poisson-like randomness, were modeled by input sharing and Poisson spike trains. As illustrated by the toy example in Figure 1 B, there are 10 cortical neurons and 4 MSNs. Each cortical neuron generates a random Poisson spike train with certain firing rate and send the signal to downstream striatal neurons. For neurons in each group of striatal neurons, blue or green, there is a distribution of input sharing due to divergent connections which would induce pair-wise correlations. We denote the setting with a parameter *W*^*in*^ = *N*_in_*/N* ^src^, within-group input sharing, where *N*_in_ is the input degree of each neuron and *N* ^src^ is the size of input source. By keeping *N*_in_ fixed (*N*_in_ = 100 in actual simulations of Figure 1 A), changing the number of *N* ^src^ changes the sharing of cortical inputs within each MSN group. In addition, the input sources are overlapped for the two striatal groups, which could range from none to totality. We denote such overlapping with a parameter *B*^*in*^ = *N* ^ol^*/N* ^src^, between-group input sharing, where *N* ^ol^ is the number of overlapped input neurons and *N* ^src^ is the size of each input source. Similarly, changing *N* ^ol^ for a given *N* ^src^ changes the sharing of cortical inputs across MSN groups. As for FSIs in Figure 1 A, they receive all inputs from both cortical sources.

### Stimulus-evoked input

To mimic stimulus-evoked response, cortical inputs were correlated by convergent connections such that two MSNs receive inputs from the same cortical neurons with different sharing probability (see section Input model). At evoked state, the average firing rates of MSN (around 5 Hz) and FSI (around 17 Hz) increases. To investigate across-trial variability, we assumed only one cortical source, hence no differentiation of groups in the striatum. We change the within-group input sharing (*W*^*in*^) as defined in section Input model from 0.01 to 0.5 to model the feed-forward sharing from cortex. For correlation transfer, the MSNs were differentiated into two groups by receiving inputs from overlapped but not identical cortical neurons. The overlapping ratio of inputs, between-group input sharing (*B*^*in*^), ranged from 0.1 to 0.9 while the within-group input sharing ranged from 0.01 to 0.5 as before.

### Statistical Analysis

#### Across-trial variability

Across-trial variability reflects the consistency of population activity in response to the same stimulus given noisy background across different trials. We quantified it as the across-trial Fano factor averaged along time. The Fano factor was calculated for each time bin (5ms) as the variance of population spike counts dividing the mean of population spike counts within the 5ms binning window across 10 trials 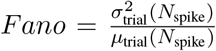 Such calculation was done for the last 2 s of the simulation and averaged as the measurement for across-trial variability. Note this metric reflects the temporally precised population variability across trials.

#### Correlation transfer

Correlation transfer denotes the transformation from input sharing (*W*^*in*^, *B*^*in*^) (see Input model and Figure 1 C) to output correlations (*W*_*out*_, *B*_*out*_). The output correlations were measured in the following way: first, all MSNs were subdivided into ensembles of size 50; then the rate trajectory of each ensemble was calculated with binning size 5 ms. The average pair-wise correlation coefficient between the rate trajectories of ensembles within each group denotes the within-group correlation, *W*_*out*_, and the average for pair of ensembles from different groups denotes the between-group correlation, *B*_*out*_.

#### Simulation and data analysis tools

The simulations were performed using the NEST simulator [Jordan et al., 2019]. Differential equations were integrated using a fixed time step of 0.1 ms. The analysis of simulated network activity was done using customized code written in Python. The results were visualized using Matplotlib.

## Data availability

Processed simulated data is shared on github https://github.com/michaelglh/FSINet.git.

## Code availability

The code for simulation and plotting is shared on github https://github.com/michaelglh/FSINet.git.

## Results

To understand the effect of highly shared FFI, we simulated the simplified cortical-striatal circuit as shown in Figure 1 and systematically varied the number of FSIs while keeping the connection in-degrees and neuron properties fixed. Here, we are interested in how the sharing of FFIs may affect the across-trial variability and the correlation structure in the evoked activity of the striatal MSNs.

### Across-trial variability

To ask whether shared FFI can affect across-trial variability, we tuned the striatal network in a baseline state with weak correlations and low firing rate [Peters et al., 2021], and injected an extra input mimicking evoked activity coming from the cortical axons (see Methods, Figure 1 A). The average firing rate of MSNs during evoked activity was ≈ 5 Hz, however, there were short bursts when MSNs spike at close to 10 Hz (Figure 2 A,B, bottom row) consistent with experimental data [Peters et al., 2021].

**Figure 2:**
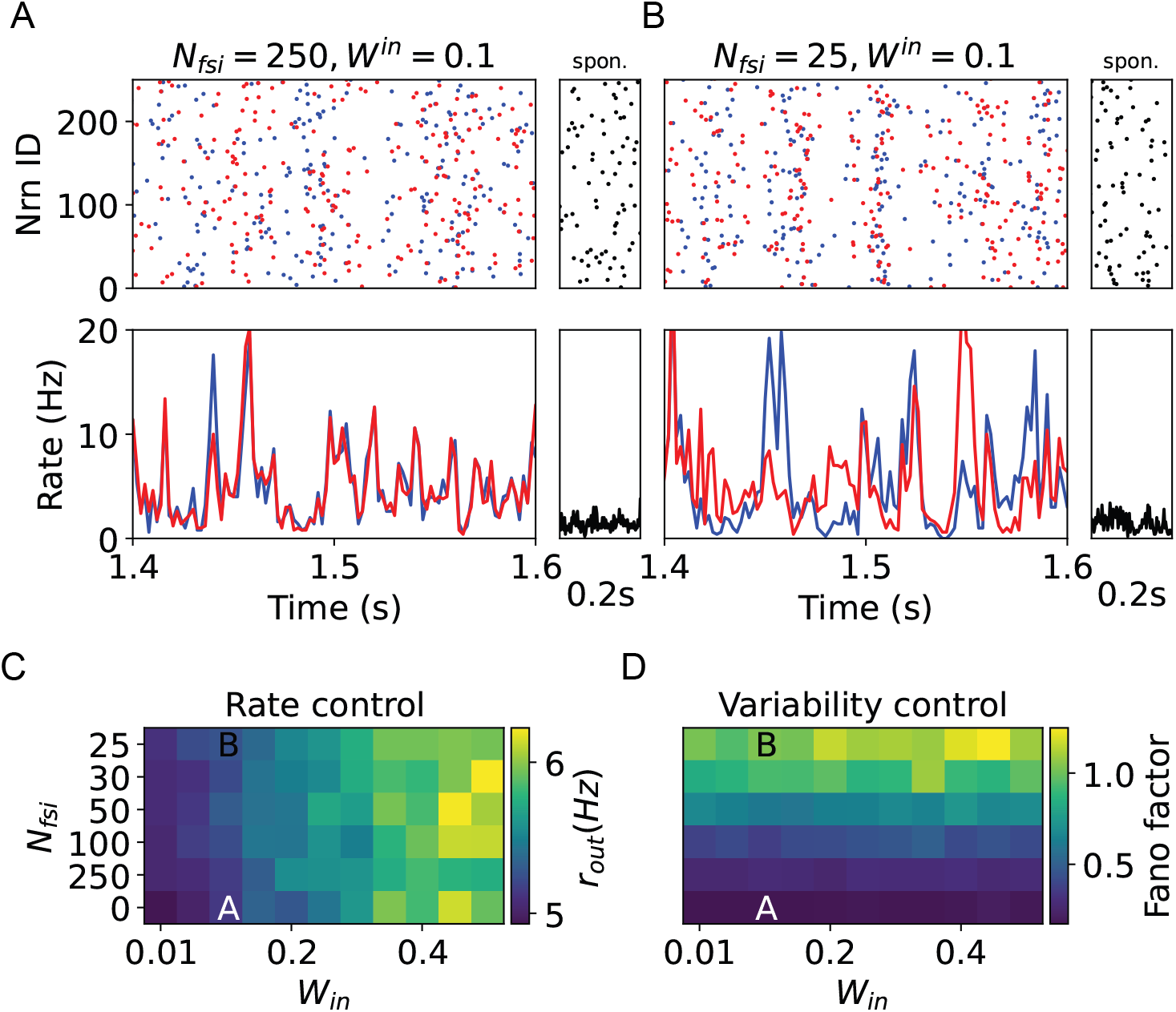
across-trial variability of MSN activities (**A**) Top: Raster plot (2 different trials) of MSN activities with *N*_fsi_ = 250, *W* ^in^ = 0.1. MSNs are stimulated with cortical inputs both by direct feedforward excitation(FFE) and FFI via FSIs. Bottom: Rate trajectories of target MSNs. The small panel shows spontaneous activity without any cortical input for 0.2s. (**B**) The rest is same as **B** but for *N*_fsi_ = 25, *W* ^in^ = 0.1. (**C**) The color denotes the average population firing rates of MSNs across trials given different FFE sharing as input source overlapping (*W* ^in^) and FFI sharing as the number of FSIs (*N*_fsi_). The labels A and B denote the two settings shown in **A** and **B**. (**D**) Same as **C** but for Fano factor.

We systematically varied the number of FSIs and cortical neurons, i.e., sharing of feedforward inhibition and feedforward excitation, and measured the across-trial mean and variance of the evoked responses. We kept the in-degree of FSI-MSN projections to maintain a relatively constant level FFI (conserved in average) hence fixed response rate of MSNs (Figure 2 C). We reduced the background inputs when *N*_fsi_ = 0 to maintain MSNs rates comparable to the case when *N*_*fsi*_ ≠ 0. Within the parameter range of *W* ^in^ and *N*_fsi_ we chose (see Stimulus-evoked input), the MSN activities could exhibit either stable Figure 2 A or variable responses Figure 2 B across trials.

We found that across-trial variability, increased with increasing sharing of FFI (fewer number of FSIs) as shown in Figure 2 D. Such effect is non-trivial as the firing rates is mostly controlled by excitatory correlation and not related to the Fano factor (compare Figure 2 C and D). The increased variability is a consequence of shared FSIs. When FSIs are shared, background activity (which is variables across trials and is assumed to be Poissonian) creates larger fluctuations in MSNs’ subthreshold membrane potentials. That is, shared FSIs create larger background noise, resulting in high across-trial variability. This effect of FSIs is further exacerbated by the fact that typical FSI inputs are on soma and therefore induce very strong inhibitory postsynaptic potentials (IPSP).

Recently, Owen et al. [2018] showed that opto-genetic inhibition of FSIs resulted in high Fano factor of MSNs’ firing rates (measured across trials). This type of variability is different from the across-trial variability we have estimated above. We considered a specific case where MSNs received stimulus evoked input in the presence and absence of FSIs. Moreover, we measured the instantaneous firing rates of the whole population. By contrast, Owen et al. [2018] measured across-trial variability of total firing rate (without any external input) of individual neurons.

### Correlation transfer attractor

It is well known that shared inputs (both excitatory or inhibitory) result in correlated activity among neurons. To avoid excessive correlations, shared input must be balanced by another input typically with opposite sign [Renart et al., 2010]. In the striatum, shared FFI can induce synchrony which can be countered by recurrent inhibition among MSNs [Yim et al., 2011]. But it is unclear how FSIs and MSNs interact when the input to the striatum is itself correlated.

To address this question, we consider that there are two groups of MSNs which received cortical inputs (see Figure 1B). This setting is used to mimic action-selection in the striatal circuit. In this simplified circuit, we do not make any assumption about the separation of D1- and D2-MSNs. However, the circuit is just sufficient to understand the role FSIs may play in transfer of correlations. The inputs to the two MSNs groups were parameterized by their mean rate, correlation within (*W*^*in*^) and between (*B*^*in*^) the inputs to either of the MSN groups (see Figure 1B). As in the previous section, here also we systematically varied the number of FSIs, *W*^*in*^, and *B*^*in*^ while maintaining the response rates of MSNs (including *N*_fsi_ = 0 without FFI). To characterize the response, we measured average the pair-wise correlation of spiking activities within each MSN group (*W*_*out*_) and between the two MSN groups (*B*_*out*_).

To compare the correlation transfer, we set the following baseline: let *N*_fsi_ = 0 such that MSNs do not get any feedforward inhibition, but they are mutually connected with recurrent inhibition. We varied input correlations (see Methods) to obtain different levels of *W*_*out*_ and *B*_*out*_. Even with the same input parameters, output correlations were variable across trials in the baseline (Figure 3 A). We clustered the output space into 25 small clusters based on measured *W*_*out*_ and *B*_*out*_ (Figure 3 A). Next, we introduced different numbers of FSIs and measured the corresponding output correlations. To visualize the effect of FSIs on correlation transfer, we plotted arrows which started from the mean *W*_*out*_ and *B*_*out*_ of baseline (see Figure 3 A) and terminated at the mean *W*_*out*_ and *B*_*out*_ in the presence of FSIs (Figure 3 B) for the 25 clusters.

**Figure 3:**
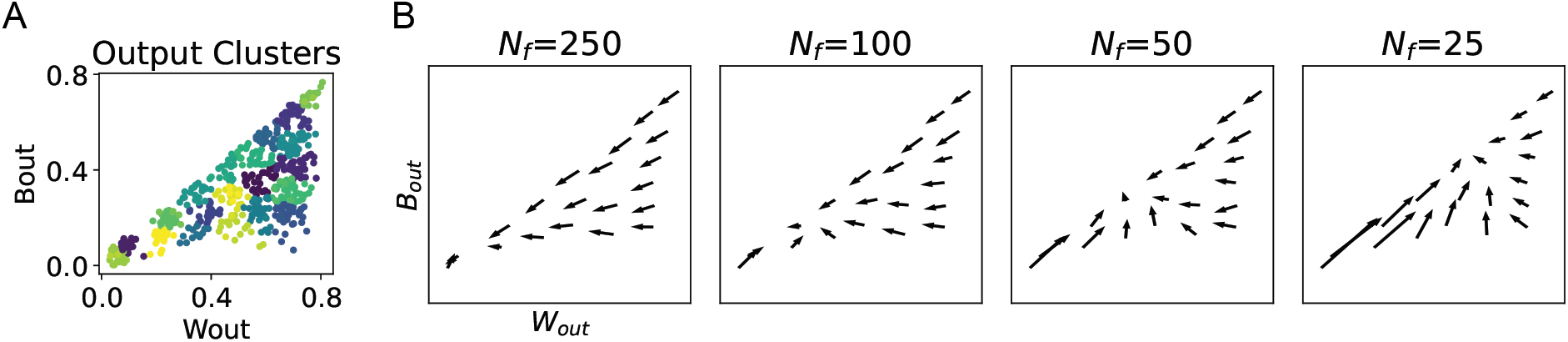
Non-linear correlation transfer in striatum. **(A)** Output correlation of MSN activities, where the x-axis is the spike correlation within the MSN group while the y-axis is the spike correlation between two MSN groups. The simulations are grouped into 25 clusters with K-means. **(B)** Each subplot compares the correlation of MSN activities between the control case (no FSI) and the studied case (certain number of FSIs). For each cluster in A, the starting points of the arrows in **B** denote the average output correlation for the control case (i.e., no FSIs), and the ending points denote the average for the studied case (*N*_f_> 0). Across columns are settings with different number of FSIs from (*N*_fsi_ = 250 to *N*_fsi_ = 25.

We found that when the network had many FSIs (10% sharing of FSIs by MSNs), MSN activities were always de-correlated within and between groups (Figure 3 B, *N*_fsi_ = 250). However, when FSIs were highly shared (60% sharing of FSIs by MSNs) as is the case in the striatum [Cinotti and Humphries, 2022], FSIs enhanced the correlation when cortical inputs were weakly correlated. In contrast, when cortical inputs were strongly correlated the striatal output was de-correlated (Figure 3 B, *N*_fsi_ = 25). Interestingly, in the right bottom region when *W*_out_ was large and *B*_out_ was small, correlations within groups were reduced while correlations between groups were increased. Therefore, in our local circuit of the striatum, the cortical differentiation of groups was weakened at striatal output. These results suggest that highly shared feedforward inhibition on top of recurrent inhibition created a *correlation attractor* in the striatum (Figure 3 B). The location of the correlation attractor monotonically moved towards zero correlations as we increased the number of FSIs (or reduced sharing of feedforward inhibition). In the previous subsection, we showed that a high degree of sharing of FSIs results in higher variability. However, sharing of FSIs did not affect the across-trial variability of output correlations.

## Discussion

Striatum is a neural network in which a very small number (only about 1%) of fast-spiking interneurons are responsible for providing feedforward inhibition to 95% MSNs. The morphology of FSIs is such that they can target a larger number of MSNs. Thus, unlike in the neocortex, feedforward inhibition in the striatum is highly shared among MSNs. It is already known that such sharing of FSIs can induce synchrony in the striatum, which can be countered by the recurrent inhibition of MSNs [Yim et al., 2011; Gittis et al., 2011]. We used a computational model to reveal two more consequences of such connectivity of FSIs: we show that shared FSIs (1) generate high across-trial variability, and (2) provide a correlation transfer attractor.

We have assumed that during evoked state MSNs fire at a relatively low average firing rate but for short periods their firing rate go up top 10 Hz. The evoked spike rate and pattern indeed depends on the task and striatal sub-region. However, our results about variability and correlation transfer are not dependent on actual operating firing rate. Even when the firing rate is fixed for MSNs, the across-trial variability of FSIs are inevitably transmitted to MSNs and sharing of FFI provides a strong modulation over correlation.

Our model predicts that given high sharing of FSI inputs, MSNs should show both high variability and synchrony. However, there are only few reports of task-related increase in synchrony among MSNs [Adler et al., 2013]. There are several possibilities how, despite sharing of FSI inputs MSNs, can maintain low synchrony. First and foremost, recurrent inhibition among MSNs can weaken the synchrony [Yim et al., 2011]. However, this effect is likely to be small given the weak recurrent inhibition among MSNs [Cinotti and Humphries, 2022]. There is another potent mechanism that can desynchronize MSNs. Depending on the reversal potential, GABA input may also be excitatory [Abed Zadeh et al., 2022]. Therefore, a strong decorrelation may occur if different MSNs show a wide range of mean membrane potential such that FSIs’ inputs appears excitatory on some MSNs. It is also possible that neuromodulators can weaken some of the FSI and thereby reduce the effective sharing of FSI inputs [Do et al., 2013].

Here we have ignored any connectivity among FSIs. Experimental data suggests that FSIs may be recurrently connected via chemical or gap-junctions. Such recurrent connections will likely increase the synchrony among FSIs [Hjorth et al., 2009; McKeon et al., 2022]. Our results show that FSIs correlated the output even when there is no synchrony. So when FSIs are themselves correlated, they may induce more synchrony and across-trial variability.

### Functional consequence

Across-trial variability can form the basis of exploration, however excessive across-trial may impair behavior. Indeed, behavioral disorders such as autism spectrum are accompanied by increased across-trial variability [Dinstein et al., 2012]. Our results suggest that FSIs control the across-trial variability. We speculate that FSIs may be useful during learning as high across-trial variability may form the basis of exploration and while performing learned tasks they may be functionally dis-engaged from MSNs to minimize across-trial variability. Consistent with this idea, optical inhibition of FSIs did not affect the performance of learned motor behavior [Owen et al., 2018] and FSIs do not provide continuous inhibition on MSNs’ firing rate [Gage et al., 2010]. The mechanism by which FSIs may be functionally disengaged from the network are not obvious at the moment.

In the absence of recurrent excitation, inhibition among MSNs can only de-correlate the activity. In such a network, FSIs provides a mechanism to boost correlations in the striatum. That is, with FSIs, striatum can also be able to bi-directionally modulate correlations. Why correlations among MSNs are needed is open to speculation, but it is easy to conceive that synchrony among MSNs, can enhance the effect of striatal activity on firing in globus pallidus (GP). As in Kuhn et al. [2003], when firing rate is fixed, the temporal correlation of input spike trains non-linearly modulates output firing rates. In our case, the *correlation attractor* possibly maintains the correlation level in a more stable range to deal with unlearned cortical representations. However, strong modulation of correlation is accompanied by high across-trial variability. The trade-off between correlation control and across-trial variability are manifested both in action control and action learning.

FSI connectivity to MSNs is increased and recurrent connectivity between MSNs is decreased in Parkinson’s disease (PD) [Gittis and Kreitzer, 2012]. According to our model, such increase of FFI and decrease of recurrent inhibition will further increase the correlation of the transfer attractor. This is consistent with increased synchrony in the striatal activity in PD [Costa et al., 2006]. In contrast to ablation of FSIs [Owen et al., 2018], it is possible that all signals are passed downstream due to strong correlations in the PD case, leading to improper action-selection. On top of it, the across-trial variability is further increased to reduce the reliability of action selection.

During action learning, Owen et al. [2018] showed that FSIs control the bursting probability of MSNs. This may mean that FSIs modulate the synaptic plasticity for MSNs. Because synaptic plasticity depends on the temporal structure of spikes [Feldman, 2012; Toyoizumi et al., 2005], the FSIs’ modulation over correlation could affect not only the level of plasticity but the structural plasticity of cortico-striatal projections. Such structural learning may be underlay by local selectivity of FSIs [Gage et al., 2010].

### Experimental verification of our results

We predict that shared FSI inputs can synchronize the MSN activity and enhance the across-trial variability. We cannot test these predictions by simply silencing the FSIs because our predictions rely on the sharing of FSIs input (or correlated FFI). Moreover, silencing will also lead to removal of inhibition and increase MSNs firing rate [Owen et al., 2018]. However, our model predicts that artificially increasing the firing rate of FSIs will further increase the variability and correlations in MSN spiking activity. Another way to verify our results is to patch two MSNs. We predict that there will be subthreshold synchrony in their membrane potential. Moreover, if we decrease the mean membrane potential of one neuron such that FSI inputs evoke a depolarization, the subthreshold synchrony between the two neurons should decrease. To the best of our knowledge, simultaneous dual patching has not been done in the striatum of awake animals however, this was already done in anesthetized animals [Stern et al., 1998]. In Parkinson’s disease, there is an increase in both the synchrony among MSNs and FSI connectivity to the MSNs [Gittis et al., 2011]. This observation is consistent with our model.

### Functional role of small neural populations

In many brain regions, some cell types exist that comprise only a small fraction of the whole population, e.g. different types of interneurons in the striatum and several cell types in the hypothalamus [Chen et al., 2017]. Such rare cell-types can affect the network activity and function in two ways: (1) by connecting in a cell-type fashion to form ensembles, and (2) by connecting with all cell types. In the later case, irrespective of whether they form excitatory or inhibitory synapses, our work suggests that such rare cell-types should affect across-trial variability and induce synchrony. The function of these aspects of the network activity will of course be contingent on the brain region.

## Acknowledgments

We thank Dr. Jens Hjerling-Leffler, Dr. Pascal Helson, Dr. Emil Waernberg, Movitz Lenninger and Hauke Wernecke for helpful discussions. This work was funded in parts by Swedish Research Council (VR), StratNeuro, Digital Futures and Inst. of Advanced Studies (University of Strasbourg, France) Fellowship grants.

